# Subtle differences in the pathogenicity of SARS-CoV-2 variants of concern B.1.1.7 and B.1.351 in rhesus macaques

**DOI:** 10.1101/2021.05.07.443115

**Authors:** Vincent J. Munster, Meaghan Flagg, Manmeet Singh, Brandi N. Williamson, Friederike Feldmann, Lizzette Pérez-Pérez, Beniah Brumbaugh, Myndi G. Holbrook, Danielle R. Adney, Atsushi Okumura, Patrick W. Hanley, Brian J. Smith, Jamie Lovaglio, Sarah L. Anzick, Craig Martens, Neeltje van Doremalen, Greg Saturday, Emmie de Wit

**Affiliations:** Laboratory of Virology, Division of Intramural Research, National Institute of Allergy and Infectious Diseases, National Institutes of Health, Hamilton, MT, United States of America; Rocky Mountain Veterinary Branch and, Division of Intramural Research, National Institute of Allergy and Infectious Diseases, National Institutes of Health, Hamilton, MT, United States of America; Research Technologies Branch, Division of Intramural Research, National Institute of Allergy and Infectious Diseases, National Institutes of Health, Hamilton, MT, United States of America

## Abstract

The emergence of several SARS-CoV-2 variants has caused global concerns about increased transmissibility, increased pathogenicity, and decreased efficacy of medical countermeasures. Animal models can be used to assess phenotypical changes in the absence of confounding factors that affect observed pathogenicity and transmissibility data in the human population. Here, we studied the pathogenicity of variants of concern (VOC) B.1.1.7 and B.1.351 in rhesus macaques and compared it to a recent clade B.1 SARS-CoV-2 isolate containing the D614G substitution in the spike protein. The B.1.1.7 VOC behaved similarly to the D614G with respect to clinical disease, virus shedding and virus replication in the respiratory tract. Inoculation with the B.1.351 isolate resulted in lower clinical scores in rhesus macaques that correlated with lower virus titers in the lungs, less severe histologic lung lesions and less viral antigen detected in the lungs. We observed differences in the local innate immune response to infection. In bronchoalveolar lavages, cytokines and chemokines were upregulated on day 4 in animals inoculated with D614G and B.1.1.7 but not in those inoculated with B.1.351. In nasal samples, we did not detect upregulation of cytokines and chemokines in D614G or B.1.351-inoculated animals. However, cytokines and chemokines were upregulated in the noses of B.1.1.7-inoculated animals. Taken together, our comparative pathogenicity study suggests that ongoing circulation under diverse evolutionary pressure favors transmissibility and immune evasion rather than an increase in intrinsic pathogenicity.

## Introduction

As the COVID-19 pandemic continues, so does the evolution of SARS-CoV-2 in the still relatively susceptible global population. The detection of the B.1.1.7 variant in the UK prompted the World Health Organization to institute a classification system for new variants, with Variants of Concern (VOC) displaying increased transmissibility or detrimental changes in COVID-19 epidemiology; increased virulence or a change in clinical disease presentation; or decreased effectiveness of public health and social measures, or available diagnostics, vaccines, or therapeutics (*1*). B.1.1.7 and B.1.351 were the first variants to be designated VOC. After initial detection, both variants spread to many other countries, fueling concerns about their transmissibility and the efficacy of vaccines. As more epidemiological data on human cases of B.1.1.7 infection become available, it appears that this variant transmits more efficiently than the variants it displaced (*2-6*), possibly due to increased virus shedding (*7*). Whether there is an increased disease severity associated with B.1.1.7 infection remains unclear (*4, 8-11*). Data on infection in humans with B.1.351 are not yet widely available, and although its rapid spread suggests increased transmissibility, there are no data to confirm this (*12-14*). Similar to B.1.1.7, it is not clear whether B.1.351 infections are associated with a change in pathogenicity compared to the other circulating SARS-CoV-2 viruses (*12-14*).

Epidemiological data on increased transmissibility and disease severity can be confounded by changes in human behavior and an increased number of cases affecting patient care. Although animal models do not completely recapitulate human disease, they are useful tools to assess several phenotypical changes such as virus transmission and pathogenesis in the absence of confounding factors. Several studies of the B.1.1.7 and B.1.351 variant in hamsters have been completed (*15-17*). However, since the changes in the spike receptor binding domain (RBD) present in these variants could affect binding to the human versus hamster ACE2 differently, replication kinetics of B.1.1.7 and B.1.351 may differ between hamsters and humans. The rhesus macaque is a widely used model of SARS-CoV-2 infection (*18*), and the amino acids in ACE2 that are essential for binding to SARS-CoV-2 spike are identical between humans and rhesus macaques (*19*). Therefore, we studied the pathogenesis of B.1.1.7 and B.1.351 in rhesus macaques and compared it to a recent SARS-CoV-2 isolate containing the D614G substitution in spike that rapidly became dominant globally in March 2020 (*20*) due to its’ increased transmissibility (*21, 22*). We found that B.1.351 was slightly less pathogenic than the other two variants, but there were no differences in virus shedding between the variants that could explain increased transmissibility.

## Methods

### Ethics and biosafety statement

All animal experiments were approved by the Institutional Animal Care and Use Committee of Rocky Mountain Laboratories, NIH and carried out in an Association for Assessment and Accreditation of Laboratory Animal Care (AAALAC) International accredited facility, according to the institution’s guidelines for animal use, following the guidelines and basic principles in the NIH Guide for the Care and Use of Laboratory Animals, the Animal Welfare Act, United States Department of Agriculture and the United States Public Health Service Policy on Humane Care and Use of Laboratory Animals. Rhesus macaques were housed in adjacent individual primate cages allowing social interactions, within a climate-controlled room with a fixed light-dark cycle (12-hr light/12-hr dark). Animals were monitored at least twice daily throughout the experiment. Commercial monkey chow was provided twice daily and the diet supplemented with treats, vegetables and/or fruit at least once a day. Water was available *ad libitum*. Environmental enrichment consisted of a variety of human interaction, manipulanda, commercial toys, videos, and music. The Institutional Biosafety Committee (IBC) approved work with infectious SARS-CoV-2 strains under BSL3 conditions. Sample inactivation was performed according to IBC-approved standard operating procedures for removal of specimens from high containment.

### Study design

To compare the pathogenicity and virus shedding of VOC B.1.1.7 and B.1.351, we inoculated three groups of six rhesus macaques with three different SARS-CoV-2 isolates: SARS-CoV-2/human/USA/RML-7/2020, a contemporary clade B.1 isolate containing the D614G substitution in spike, hCOV_19/England/204820464/2020, a B.1.1.7 isolate, and hCoV-19/USA/MD-HP01542/2021, a B.1.351 isolate. Each group of six animals contained one older adult animal (age 15-16 years), one adult animal (3-5 years) and four young rhesus macaques (age 2-3 years). All but one of the older animals were male. Each group of animals was housed in a separate room. The animals were inoculated intranasally via a MAD Nasal™ IN Mucosal Atomization Device (Teleflex, US; 0.5ml per nostril) and intratracheally (4 ml) with an inoculum of 4×10^5^ TCID50/ml virus dilution in sterile DMEM, resulting in a total dose of 2×10^6^ TCID50. The inocula were titered on Vero E6 cells to confirm the correct dose was administered. The animals were observed and scored daily according to a standardized scoring sheet (*23*); the same person, blinded to the isolate the animals received, assessed the animals throughout the study. The predetermined endpoint for this experiment was 6 days post inoculation (dpi). Clinical exams were performed on 0, 2, 4, 6 dpi on anaesthetized animals. On exam days, clinical parameters such as bodyweight, body temperature and respiratory rate were collected. Additionally, ventro-dorsal and right/left lateral thoracic radiographs were taken prior to any other procedures (e.g. bronchoalveolar lavage, nasal flush). Radiographs were evaluated and scored for the presence of pulmonary infiltrates by two board-certified clinical veterinarians according to a standard scoring system. Briefly, each lung lobe (upper left, middle left, lower left, upper right, middle right, lower right) was scored individually based on the following criteria: 0 = normal examination; 1 = mild interstitial pulmonary infiltrates; 2 = moderate interstitial pulmonary infiltrates, perhaps with partial cardiac border effacement and small areas of pulmonary consolidation (alveolar patterns and air bronchograms); and 3 = pulmonary consolidation as the primary lung pathology, seen as a progression from grade 2 lung pathology. At study completion, thoracic radiograph findings are reported as a single radiograph score for each animal on each exam day. To obtain this score, the scores assigned to each of the six lung lobes are added together and recorded as the radiograph score for each animal on each exam day. Scores may range from 0 to 18 for each animal on each exam day. Blood as well as nasal, throat, and rectal swabs were collected during all clinical exams. Nasosorption swabs were collected from the nostril that was not used for nasal swabbing; nasosorption swabs were inserted in the nostril and kept in place for 1 minute.

Nasosorption swabs were collected in 300µl PBS containing 1% BSA and 0.4% Tween-20 and vortexed for 30 seconds. The swab and liquid were then placed on a spin filter (Agilent, 5185-5990) and centrifuged at 16,000 rpm for 20 minutes. Filtered liquid was aliquoted and stored at −80°C. Additionally, on 2 and 4 dpi animals were intubated and bronchoalveolar lavages (BAL) were performed using 10ml sterile saline and a second sample was collected in the same location using a cytology brush (bronchial cytology brush, BCB). The brush was then placed in 1ml DMEM containing 50 U/ml penicillin and 50 μg/ml streptomycin. On 6 dpi, all animals were euthanized; after euthanasia, necropsies were performed and tissue samples were collected. Histopathological analysis of tissue slides was performed by a board-certified veterinary pathologist blinded to the group assignment of the animals.

### Virus and cells

SARS-CoV-2/human/USA/RML-7/2020 (GenBank: MW127503.1; designated D614G throughout the manuscript) was obtained from a nasopharyngeal swab obtained on July 19, 2020. Two passages of the original swab were performed in VeroE6 cells. The virus stock used was 100% identical to the deposited Genbank sequence. hCoV-19/England/204820464/2020 (GISAID: EPI_ISL_683466; designated B.1.1.7 throughout the manuscript) was obtained from Public Health Agency England via BEI Resources. The obtained passage 2 material was propagated once in VeroE6 cells. Sequencing confirmed the presence of three SNPs in this stock: nsp6 D156G (present in 14% of all reads), nsp6 L257F (18%) and nsp7 V11I (13%). hCoV-19/USA/MD-HP01542/2021 (GISAID: EPI_ISL_890360; designated B.1.351 throughout the manuscript) was obtained from Dr. Andrew Pekosz at the Johns Hopkins University Bloomberg School of Public Health. The obtained passage 2 material was propagated once in Vero E6 cells. Sequencing confirmed the presence of two SNPs in this stock: nsp5 P252L (17%) and nsp6 L257F (57%). The D614G, B.1.1.7 and B.1.351 stocks contained similar ratios of SARS-CoV-2 E gene copies to infectious virus, i.e. 1.3×10^4^, 1.6×10^4^ and 1.5×10^4^ RNA copies/TCID50, respectively.

VeroE6 cells were maintained in DMEM supplemented with 10% fetal bovine serum, 1 mM L-glutamine, 50 U/ml penicillin and 50 μg/ml streptomycin; SARS-CoV-2 stocks were grown in the same medium, except with 2% fetal bovine serum.

### Quantitative PCR

RNA was extracted from swabs and BAL using the QiaAmp Viral RNA kit (Qiagen) according to the manufacturer’s instructions. Tissues (30 mg) were homogenized in RLT buffer and RNA was extracted using the RNeasy kit (Qiagen) according to the manufacturer’s instructions. For detection of genomic and subgenomic RNA, 5 µl RNA was used in a one-step real-time RT-PCR assay (*24, 25*) using the QuantiFast probe kit (Qiagen) according to instructions of the manufacturer. In each run, standard dilutions of counted RNA standards were run in parallel, to calculate copy numbers in the samples.

### Virus titration

Virus titrations were performed by end-point titration in Vero E6 cells. Tissues were homogenized in 1ml DMEM using a TissueLyser II (Qiagen) and a 5 mm stainless steel bead. Cells were inoculated with 10-fold serial dilutions of swab, BAL, BCB, or homogenized tissue samples in 96-well plates. Plates were centrifuged for 30 minutes at 1000rpm and incubated for 30 minutes at 37°C and 5% CO_2_. The inoculum was then removed and replaced with 100 µl DMEM containing 2% FBS, 50 U/ml penicillin and 50 μg/ml streptomycin. Six days after inoculation, CPE was scored and the TCID50 was calculated.

### Next generation sequencing of viral RNA

Viral RNA was extracted as described above. Kapa’s RNA HyperPrep library preparation kit (Roche Sequencing Solutions) was used to prepare sequencing libraries 10µl RNA. To facilitate multiplexing, adapter ligation was performed with the KAPA Universal Adapter and Unique Dual-Indexed Primer mixes. Samples were enriched for adapter-ligated product using KAPA HiFi HotStart Ready mix and 12 PCR amplification cycles, according to the manufacturer’s manual. Pools consisting of eight to ten sample libraries were used for hybrid-capture virus enrichment using myBaits^®^ Expert Virus SARS-CoV-2 panel and following the manufacturer’s manual, version 4.01, with eight - 13 cycles of post-capture PCR amplification (Arbor Biosciences). Purified, enriched libraries were quantified using Kapa Library Quantification kit (Roche Sequencing Solutions) and sequenced as 2 × 150 base pair reads on the Illumina NextSeq 550 instrument (Illumina).

Raw fastq reads were trimmed of Illumina adapter sequences using cutadapt version 1.12 (*26*) and then trimmed and filtered for quality using the FASTX-Toolkit (Hannon Lab). Remaining reads were mapped to the the respective hCOV_19/England/204820464/2020, hCoV-19/USA/MD-HP01542/2021, and SARS-CoV-2/human/USA/RML-7/2020 genomes using Bowtie2 version 2.2.9 with parameters --local --no-mixed -X 1500. PCR duplicates were removed using picard MarkDuplicates (Broad Institute) and variants were called using GATK HaplotypeCaller version 4.1.2.0 (*27*) with parameter-ploidy 2. Variants were filtered for QUAL > 500 and DP > 20 using bcftools.

### Histopathology

Histopathology and immunohistochemistry were performed on rhesus macaque tissues. After fixation for a minimum of 7 days in 10 % neutral-buffered formalin and embedding in paraffin, tissue sections were stained with hematoxylin and eosin (HE). Immunohistochemistry was performed using a custom-made rabbit antiserum against SARS-CoV-2 N at a 1:1000 dilution. Stained slides were analyzed by a board-certified veterinary pathologist. Histologic lesion severity was scored per lung lobe according to a standardized scoring system evaluating the presence of interstitial pneumonia, type II pneumocyte hyperplasia, edema and fibrin, and perivascular lymphoid cuffing: 0, no lesions; 1, minimal (1-10% of lobe affected); 2, mild (11-25%); 3, moderate (26-50%); 4, marked (51-75%); 5, severe (76-100%).

Presence of viral antigen was scored per lung lobe according to a standardized scoring system: 0, none; 1, rare/few; 2, scattered; 3, moderate; 4, numerous; 5, diffuse.

### Cytokine and chemokine analysis

Serum, BAL fluid and nasosorption samples were analyzed for the presence of GM-CSF, MCP-1, TNF-α, IFN-α2a, IFN-γ, IL-1RA, IL-1β, IL-6, IL-8 and IL-15 using the U-PLEX NHP Biomarker Group 1 kit (Meso Scale Diagnostics) according to the manufacturer’s instructions. Plates were read using a MESO QuickPlex SQ 120 instrument (Meso Scale Diagnostics). Samples from animals inoculated with different variants were randomized across plates to mitigate batch effects. Samples were inactivated via 2MRad gamma-irradiation prior to analysis outside of containment (*28*). For BAL samples, 0.1% Triton X-100 (Sigma Aldrich) was added immediately before running the assay to prevent clumping. Analyte concentrations and upper and lower detection limits were determined using the Discovery Workbench software (Meso Scale Diagnostics) individually for each assay plate. Samples that fell below the lower detection limit were replaced with the corresponding lower detection limit value. Data were analyzed using Python (version 3.8.5) and GraphPad Prism (version 8.2.1). To correct for baseline differences in cytokine concentrations between animals, the log2 fold change of analytes was calculated on a per-animal basis, using the 0 dpi sample as a baseline. Log2 fold changes were not calculated for BAL samples, as a 0 dpi sample was not available. Unsupervised hierarchical clustering (Ward’s) was performed on Euclidean pairwise distance matrices calculated from the mean of the log2 fold change or concentration of analytes from animals in each group at each available timepoint using SciPy (version 1.6.0) and Seaborn (version 0.11.1) packages in Python.

### Statistical analysis

Statistical analyses were performed using GraphPad Prism software version 8.2.1. A p-value of 0.05 was used as cut off for statistical significance.

### Data availability

All data included in this manuscript have been deposited in Figshare at https://figshare.com/articles/dataset/dx_doi_org_10_6084_m9_figshare_6025748/6025748. Sequences have been deposited in NCBI under BioProject Accession number PRJNA727012.

## Results

### Reduced disease in B.1.351-inoculated rhesus macaques

The pathogenicity of two VOC isolates, B.1.1.7 and B.1.351, was compared to the pathogenicity of a recent clade B.1 isolate containing the D614G substitution in the spike protein. Three groups of six rhesus macaques were inoculated intranasally and intratracheally with a total dose of 2×10^6^ TCID50 of one of the following SARS-CoV-2 isolates: SARS-CoV-2/human/USA/RML-7/2020, containing the D614G substitution in spike, hCOV_19/England/204820464/2020, a B.1.1.7 isolate, and hCoV-19/USA/MD-HP01542/2021, a B.1.351 isolate.

Clinical signs were mild in all three groups, with the main clinical signs observed being reduced appetite and changes in respiratory pattern (Fig. S1). Additionally, a clear nasal discharge was observed on 6 dpi in a single animal inoculated with D614G. Clinical scores were significantly lower in the animals inoculated with B.1.351 than B.1.1.7 and D614G on several days after inoculation (Fig. 1A). Further analysis of the clinical scores showed that this was due to a combination of differences in appetite as well as respiratory signs. The B.1.351-inoculated animals had fewer days with a reduced appetite and only 3 out of 6 animals inoculated with B.1.351 showed respiratory signs of disease at any time after inoculation, compared to 5 out of 6 for D614G and 4 out of 6 for B.1.1.7 (Fig. S1). Radiographs collected on 0, 2, 4, 6 dpi were analyzed for the presence of pulmonary infiltrates. Pulmonary infiltrates were observed in the majority of inoculated animals; however, no differences were observed between the three groups in the severity of infiltrates (Fig. 1B).

**Figure 1.**
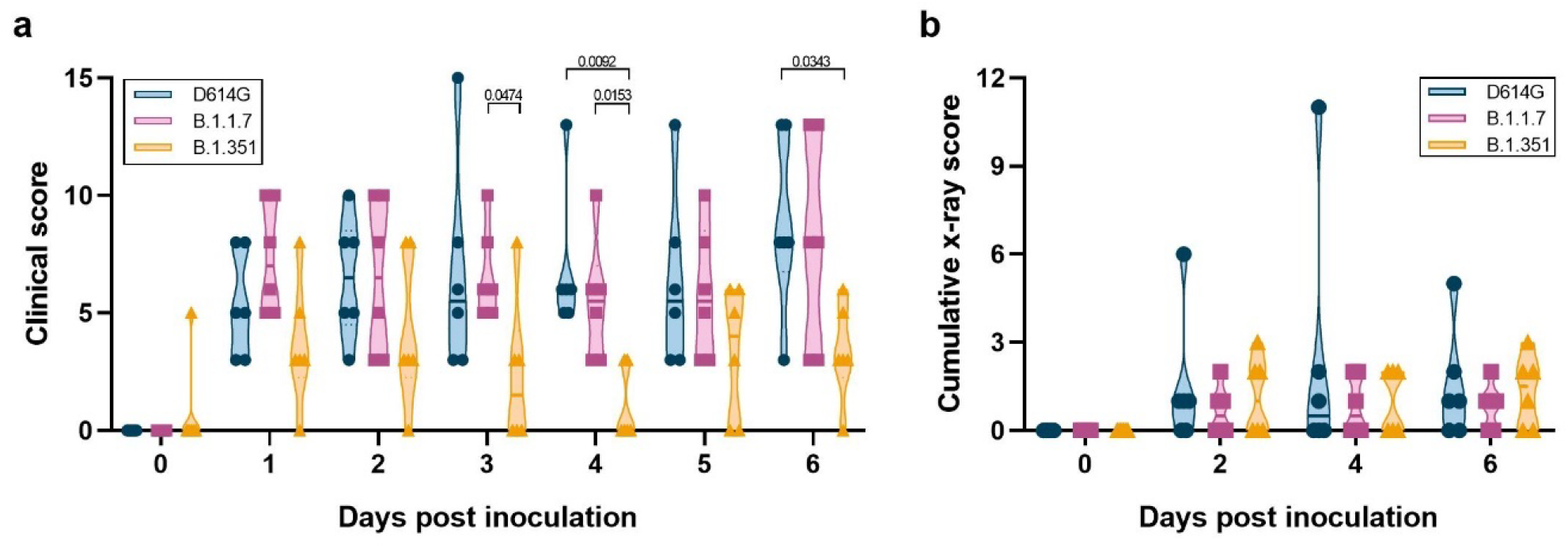
Milder disease observed in rhesus macaques inoculated with B.1.351 than with D614G or B.1.1.7. Three groups of six adult rhesus macaques were inoculated with SARS-CoV-2 variants D614G, B.1.1.7 or B.1.351. After inoculation, animals were observed for disease signs and scored according to a pre-established clinical scoring sheet (a). Ventro-dorsal and lateral radiographs were taken on clinical exam days and scored for the presence of pulmonary infiltrates by a clinical veterinarian according to a standard scoring system. Individual lobes were scored and scores per animal per day were totaled and displayed (b). Center bars indicate the mean. Statistical analysis was performed using a 2-way ANOVA with Tukey’s multiple comparisons test; p-values <0.05 are indicated.

### No difference in virus shedding between the three groups

Since increased virus shedding may lead to increased transmission efficiency, we studied virus shedding after inoculation as a proxy for transmission potential. Nose, throat and rectal swabs were collected on 2, 4, and 6 dpi and analyzed for the presence of genomic RNA (gRNA), subgenomic RNA (sgRNA; an indicator of recent virus replication (*29*)) and, in the case of nose and throat swab, infectious virus.

High amounts of gRNA, sgRNA and high virus titers were observed in nose swabs (Fig. 2A) on 2 dpi, and these slowly declined over time. No differences in virus shedding from the nose were observed between the three groups. A similar pattern of virus shedding was observed in the throat swabs (Fig. 2B), again with no differences between the three groups. As previously observed (*23, 30, 31*), virus shedding was less consistent in the rectal swabs than in nose or throat swabs, but again there were no differences between the three groups (Fig. 2C). To determine whether the total amount of virus shedding was different between the three groups, we also calculated the area under the curve for shedding of gRNA, sgRNA and infectious virus in all swabs. No statistically significant differences in total amount of shedding were detected (Fig. S2A).

**Figure 2.**
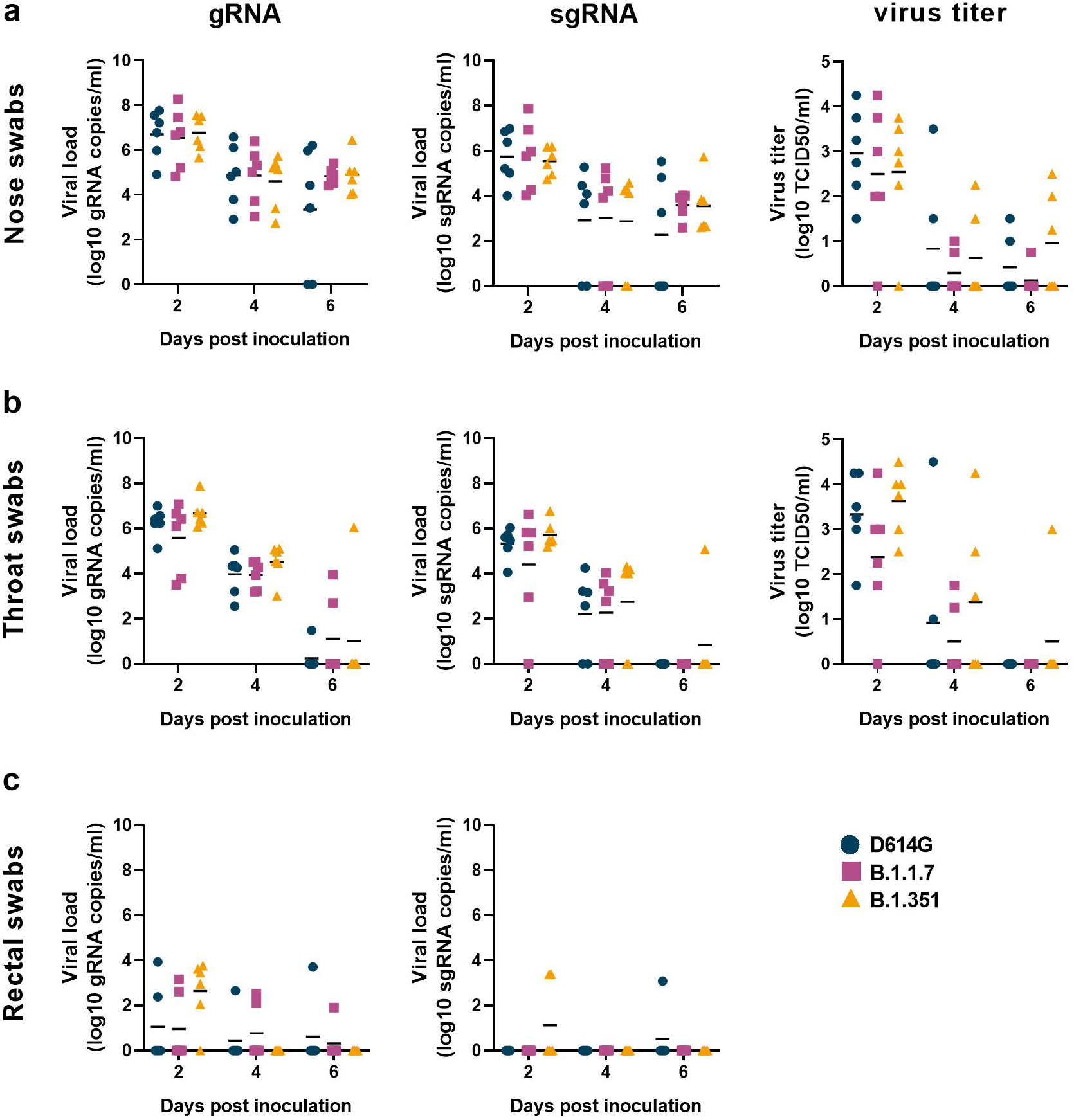
No differences in virus shedding between D614G, B.1.1.7 and B.1.351. Three groups of six adult rhesus macaques were inoculated with SARS-CoV-2 variants D614G, B.1.1.7 or B.1.351. After inoculation, clinical exams were performed during which nose (a), throat (b), and rectal (c) swabs were collected; viral loads and virus titers were measured in these samples. qRT-PCR was performed to detect genomic (left column) and subgenomic RNA (middle column), and virus titration was performed on nose and throat swabs to detect levels of infectious virus (right column) in these samples. Lines indicate the mean. Statistical analysis was performed using a 2-way ANOVA with Tukey’s multiple comparisons test; no p-values <0.05 were found.

### Similar virus replication kinetics in the airways during infection

As an indicator of virus replication in the lower respiratory tract during infection, bronchoalveolar lavages (BAL) and bronchial cytology brush (BCB) samples were collected on 2 and 4 dpi and analyzed for the presence of gRNA, sgRNA and infectious virus. Viral loads and virus titer in BAL and BCB were highest on 2 dpi and declined by 4 dpi (Fig. S2B, C). Although viral loads and virus titers were generally higher in the BAL fluid than in the BCB sample, these two samples showed very similar kinetics overall, with samples with low viral loads in BAL also having low viral loads in BCB. No statistically significant differences were detected in viral RNA or virus titer in BAL or BCB between the three variants (Fig. S2B, C).

To determine whether SNPs detected in the B.1.1.7 and B.1.351 virus stocks were stable *in vivo*, we performed Next Generation Sequencing of BAL samples collected on 2dpi, close to the peak of virus replication. No SNPs were detected in the D614G inoculum at an allelic fraction >0.1, and only one SNP was detected in the BAL sample of one animal; this was a synonymous mutation in nsp6 (Table S1). The B.1.1.7 inoculum contained 3 SNPs at >0.1 allelic fraction compared to the reference sequence; the D156G in nsp6 was detected in all animals, but at an allelic fraction <0.1 in 5 out of 6 animals. The L257F substitution in nsp6 was maintained in all animals, and the V11I substitution increased in frequency in 5 out of 6 animals (Table S1). The B.1.351 inoculum contained 2 amino acid substitutions compared to the reference sequence; of these, the P252L substitution in nsp5 was maintained in 5 of 6 animals at slightly higher percentages than in the inoculum, whereas the L257F substitution in nsp6 was maintained at levels similar to the virus inoculum in all animals (Table S1). Overall, the vast majority of amino acid substitutions present in the B.1.1.7 and B.1.351 inocula was maintained during replication in rhesus macaques, indicating that the effect of these substitutions on virus replication was most likely neutral in this host.

### Lower virus titer in the lungs of B.1.351-inoculated animals

On 6 dpi, all animals were euthanized and necropsies were performed. To determine whether the presumed increased transmission of B.1.1.7 or B.1.351 could be due to higher levels of virus replication in the nasal turbinates, we collected nasal turbinate material from three sites in the nasal turbinates and analyzed these for the presence of gRNA, sgRNA, and infectious virus (Fig. 3A). High viral loads of gRNA and sgRNA were detected in the nasal turbinates of all animals; fewer tissue samples were positive for sgRNA than gRNA, indicating virus replication had peaked before the time of sampling. There were no statistically significant differences in viral loads between the variants (Fig. 3A). Fewer nasal turbinate samples had detectable infectious virus in the group of B.1.351-inoculated animals than in those inoculated with D614G or B.1.1.7, but this difference was not statistically significant (Fig. 3A). Similar analyses were performed on samples collected from all 6 lung lobes. Again, high viral loads of gRNA and sgRNA were detected, with fewer tissue samples being positive for sgRNA than gRNA (Fig. 3B).

**Figure 3.**
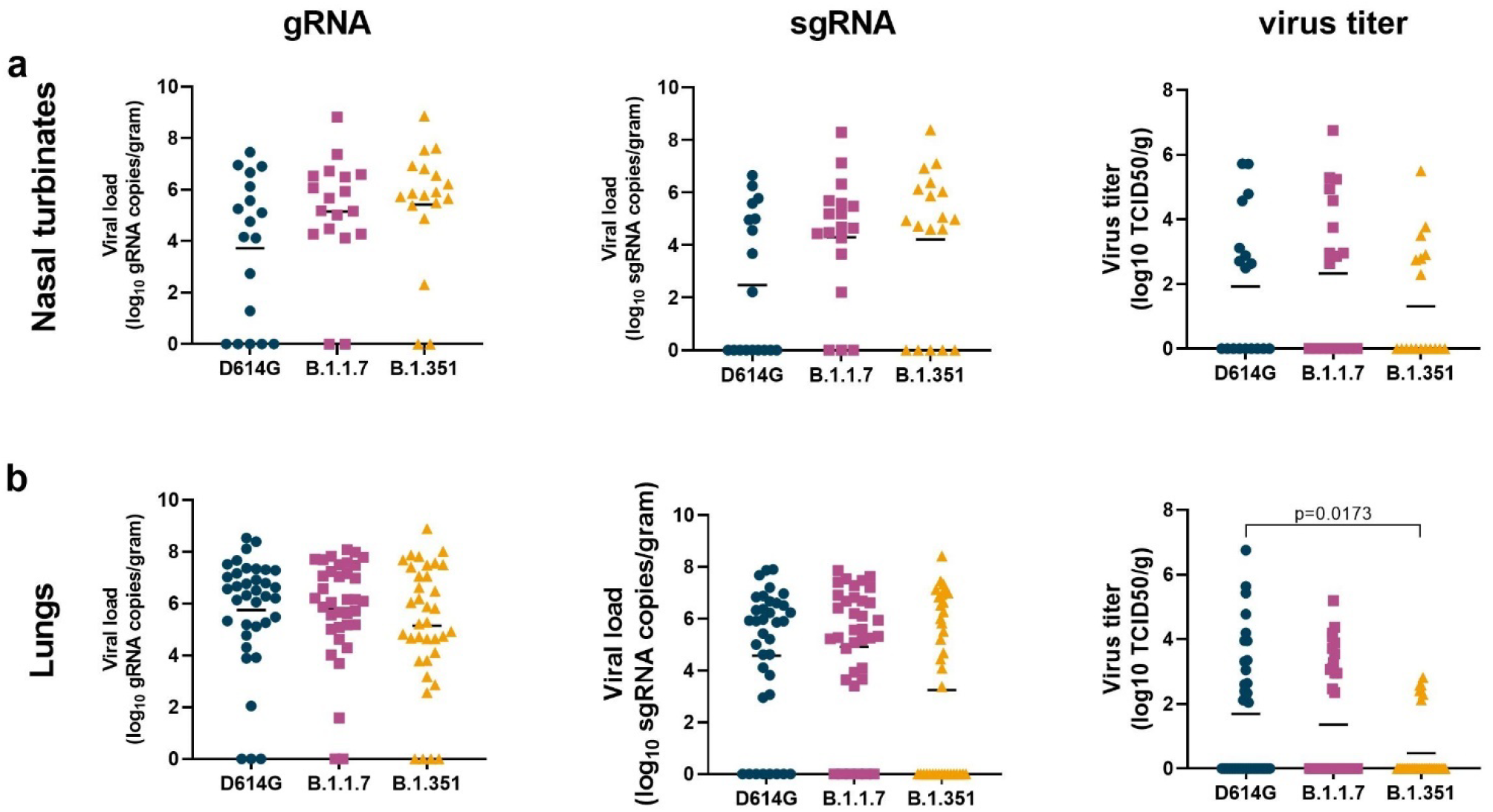
Lower virus titers in lungs, but not nasal turbinates, of B.1.351-inopculated rhesus macaques. Three groups of six adult rhesus macaques were inoculated with SARS-CoV-2 variants D614G, B.1.1.7 or B.1.351. On 6 dpi, all animals were euthanized and necropsies were performed. Samples were collected from the frontal, mid and rear nasal turbinates (a) as well as all 6 lung lobes (b) and analyzed for the presence of gRNA (left panels), sgRNA (middle panels) and virus titer (right panels). Lines indicate mean. Statistical analysis was performed using a Kruskal-Wallis test with Dunn’s multiple comparisons tests and p-values <0.05 are indicated.

Significantly lower virus titers were detected in the lungs of B.1.351-inoculated animals than in those inoculated with D614G (Fig. 3B), possibly explaining the lower clinical scores observed in the animals inoculated with B.1.351 (Fig. 1A).

Additional tissue samples were analyzed for the presence of gRNA and sgRNA only. In the nasal mucosa, gRNA levels were significantly higher in B.1.1.7-inoculated animals than in the other two groups (Fig. S3); however, this difference was not observed in sgRNA levels in the same tissues and thus probably does not represent increased virus replication in this tissue. Although B.1.1.7 appears to be detected in the various tissue samples more frequently than D614G and B.1.351, these differences were not statistically significant in individual tissues (Fig. S3).

### Less severe histologic lesions and less viral antigen detected in the lower respiratory tract of B.1.351-inoculated rhesus macaques

Lung tissues collected on 6 dpi were analyzed for the presence of histologic lesions. All inoculated animals developed some degree of pulmonary pathology, resulting in lesions typical of SARS-CoV-2 infection in rhesus macaques. On observation, lung lesions in animals inoculated with D614G were generally more severe than B.1.1.7 and B.1.351 while there were minimal differences between animals inoculated with B.1.1.7 and B.1.351 (Fig. 4A-F). Analysis of the histology scores assigned to each lung lobe of all animals indicated that lesions in D614G, B.1.1.7 and B.1.351-inoculated animals occurred on a gradient from more to less severe, respectively; these differences were statistically significantly more severe in D614G and B.1.1.7-inoculated animals than in B.1.351-inoculated animals (Fig. 4J).

**Figure 4.**
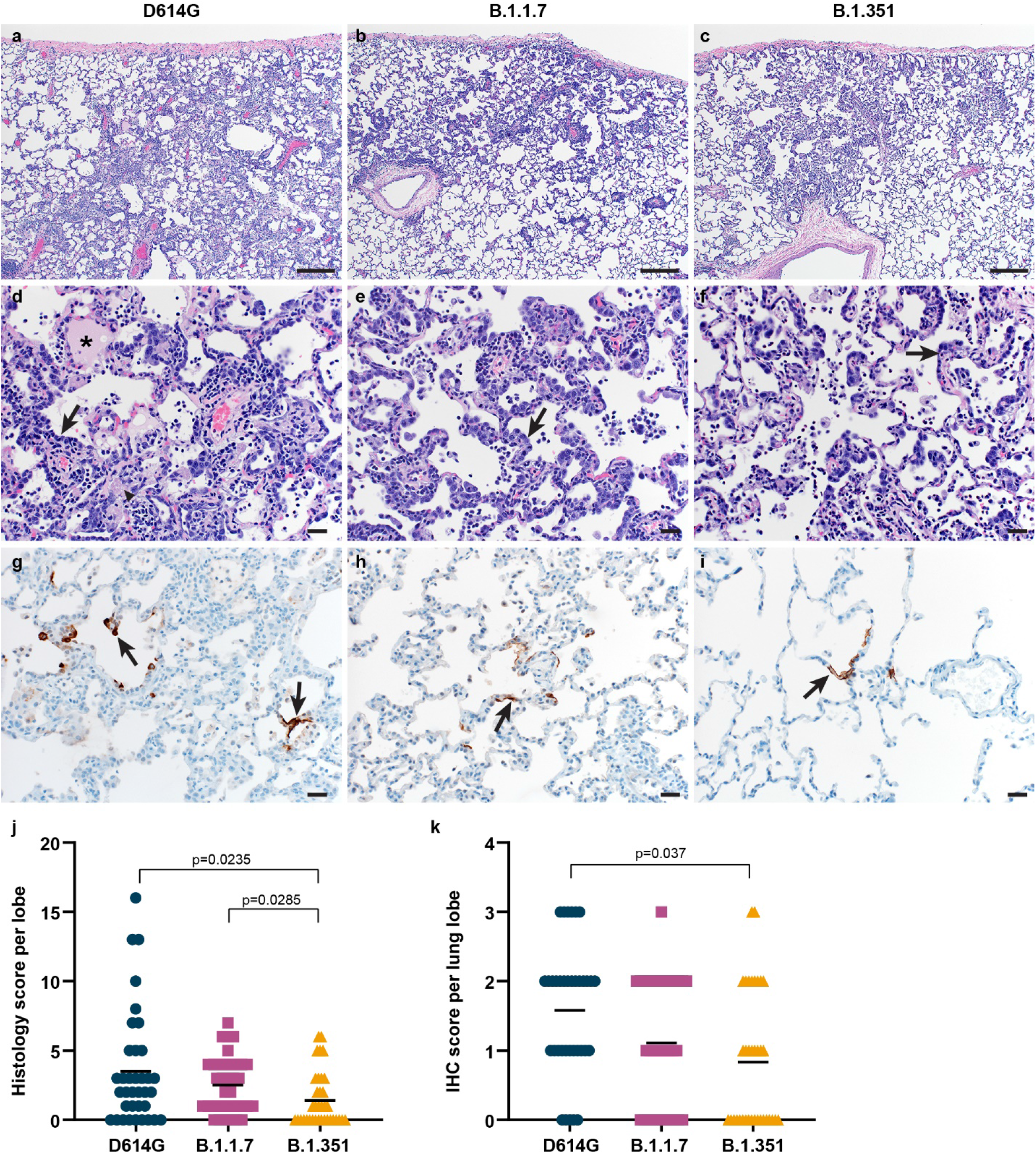
Differences in histopathological changes in lungs of rhesus macaques inoculated with D614G,B.1.1.7 or B.1.351. Three groups of six adult rhesus macaques were inoculated with SARS-CoV-2 variants D614G, B.1.1.7 or B.1.351. On 6 dpi, all animals were euthanized and necropsies were performed. Lungs were assessed for the presence of interstitial pneumonia. Moderate (a) and mild (b, c) interstitial pneumonia were observed. In moderate lesions, edema (asterisk), fibrin (arrow head), and type II pneumocyte hyperplasia (arrow) were detected (d). Type II pneumocyte hyperplasia was observed in the mild lesions (e, f). By immunohistochemistry for SARS-CoV-2 N protein, moderate amounts of antigen-positive pneumocytes (arrows) were detected in D614G-inoculated animals (g), whereas scattered numbers of antigen-positive pneumocytes (arrows) were detected in B.1.1.7- and B.1.351-inoculated animals (h, i). Size bars indicate 200µM (a-c) or 20 µM (d-i). Histological lesion severity was scored per lung lobe according to a standardized scoring system evaluating the presence of interstitial pneumonia, type II pneumocyte hyperplasia, edema and fibrin, and perivascular lymphoid cuffing (score 0-5); these values were combined per lung lobe and graphed (j). Presence of viral antigen was scored per lung lobe according to a standardized scoring system (0-5); these values were combined per lung lobe and graphed (k). Statistical analysis (j, k) was performed using a Kruskal-Wallis test with Dunn’s multiple comparisons tests and p-values <0.05 are indicated.

Immunohistochemistry was performed to detect SARS-CoV-2 antigen in lung tissue. As shown previously, viral antigen was present in type I and II pneumocytes, as well as alveolar macrophages. On observation, the animals inoculated with D614G generally had more viral antigen than those inoculated with B.1.1.7 or B.1.351 (Fig. 4G-I). Analysis of the abundance of antigen presence in each lung lobe of all animals again indicated a gradient of viral antigen abundance from D614G to B.1.1.7 and B.1.351. There was statistically significantly more antigen present in the lungs of D614G-inoculated animals than in those inoculated with B.1.351 (Fig. 4K), in line with the lower virus titers detected in the lungs of these animals.

### Subtle differences in local, but not systemic innate immune response

We analyzed serum, BAL and nasosorption samples for the presence of 10 cytokines and chemokines and found different responses in the different sample types. In serum, the immune response peaked at 2 dpi, with animals demonstrating up-regulation of IL-6, IL-15, IL-1RA, MCP-1, IFN-α2a, and TNF-α, and was largely resolved by 4 dpi (Fig. 5A). Subtle differences between variant groups were observed in the levels of up-regulation of several cytokines, but the overall responses were largely similar (Fig. S4A). Unsupervised hierarchical clustering confirmed this observation, with samples clustering by timepoint rather than variant group (Fig. 5A).

**Figure 5.**
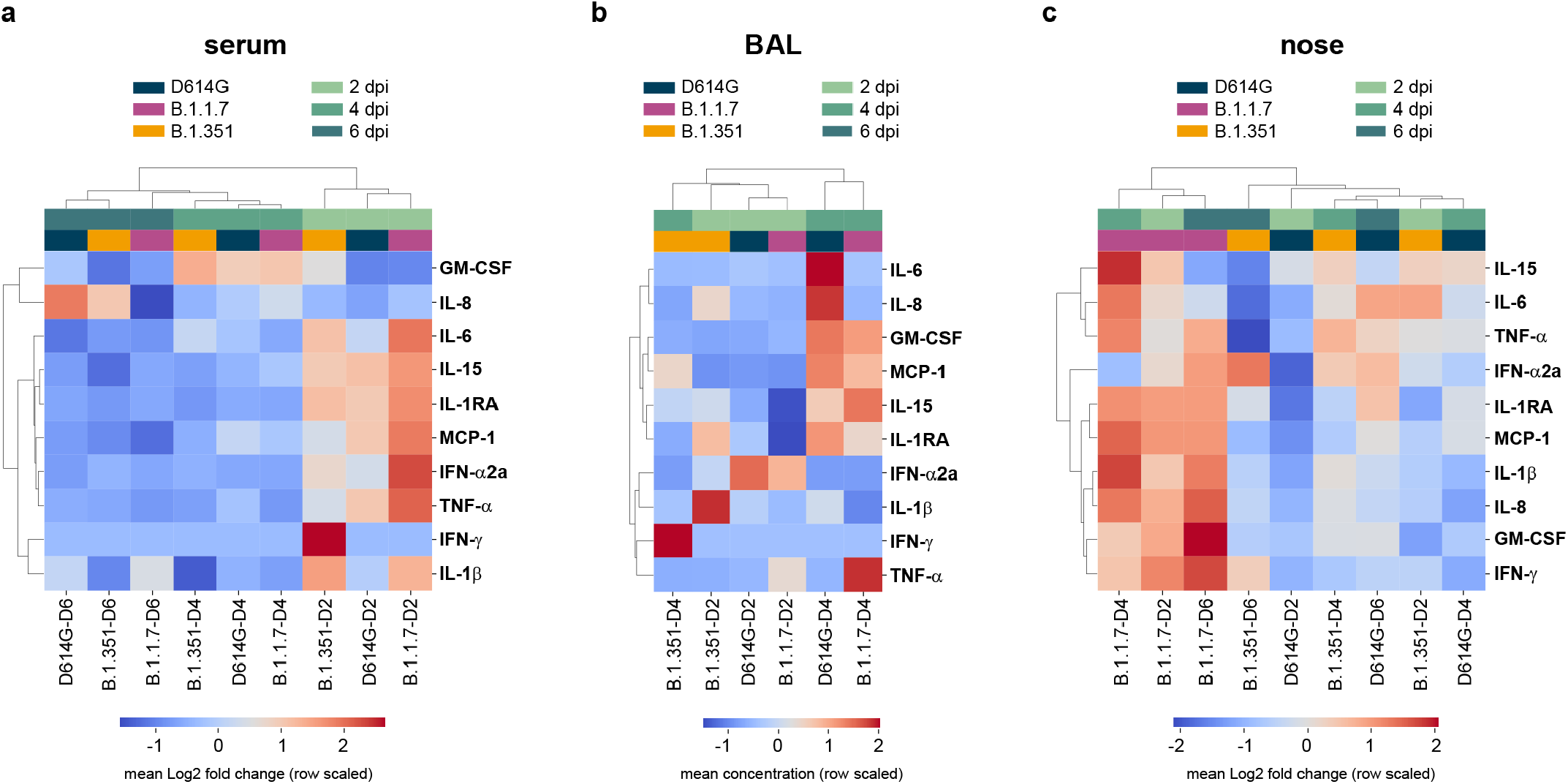
Differences in cytokine and chemokine levels in serum, BAL and nose samples of rhesus macaques inoculated with D614G, B.1.1.7 or B.1.351. Three groups of six adult rhesus macaques were inoculated with SARS-CoV-2 variants D614G, B.1.1.7 or B.1.351. The levels of 10 cytokines and chemokines were determined in serum on 0, 2, 4, and 6 dpi (a); in BAL on 2 and 4 dpi (b); and in nasal samples on 0, 2, 4, and 6 dpi (c). For serum and nasal samples, fold change in cytokine/chemokine level is shown normalized to baseline levels measured on 0 dpi. For BAL samples, concentrations are shown. Data represent the mean log2 fold change or concentration of analytes in variant groups at each timepoint, and are organized according to unsupervised hierarchical clustering.

The cytokine and chemokine response in BAL was slightly delayed compared to that in serum, with the peak response at 4 dpi. This was defined by up-regulation of GM-CSF, MCP-1, and IL-15, indicating a concerted recruitment of leukocytes (Fig. 5B, S4B). Notably, this response was absent in B.1.351-inoculated animals as indicated by these samples clustering separately from D614G- and B.1.1.7-inoculated animals (Fig. 5B). The limited immune response in the BAL of B.1.351-inoculated animals is in agreement with the reduced lung virus titers and histologic lesion severity observed in these animals (Fig. 3B, 4I-J), suggesting reduced pathogenicity in the lower respiratory tract.

In nasal samples, we observed a striking difference in the immune response in B.1.1.7-inoculated animals compared to those inoculated with D614G and B.1.351. Samples from B.1.1.7-inoculated animals at all timepoints clustered separately, and showed a unique pro-inflammatory response consisting of up-regulation of IL-6, MCP-1, IL-1β, IL-8, and GM-CSF that persisted up to 6 dpi (Fig. 5A, S4C). Interestingly, B.1.1.7-inoculated animals were the only group that did not down-regulate IFN-γ (Fig. S4C).

## Discussion

Despite reports of a potential increase in disease severity in humans (*10, 12-14*), B.1.1.7 and B.1.351 did not show increased pathogenicity in rhesus macaques. In fact, the B.1.351 isolate appears to be less pathogenic in these animals. However, the absence of increased disease in individuals does not necessarily mean an overall decrease in virulence, since increased transmission of these variants could result in an increased disease burden of these variants on the population level (*32*).

The reduced replication of B.1.351 in the lower, but not upper respiratory tract, might imply that this virus is evolving to become more like a virus of the upper respiratory tract. Likewise, the only difference observed in our study between B.1.1.7 and D614G was in the innate immune response in the upper, but not lower respiratory tract.

The virus shedding data in swabs collected from the upper respiratory tract did not directly explain the increased transmission of B.1.1.7 and B.1.351 observed in the population. However, the upregulation of innate immune responses in the nasal samples collected from B.1.1.7-inoculated animals may potentially play a role in the increased transmission of this variant. Nasal inflammatory responses such as the increased cytokine and chemokine concentrations observed here have been linked to common cold symptoms in humans (*33, 34*). Symptomatic 229E coronavirus infection in human volunteers was linked to increased plasma exudation into the nasal cavity (*35*). One potential explanation for increased transmission of B.1.1.7 could be that increased protein concentrations in nasal secretions caused by plasma exudation could stabilize the virus in respiratory droplets (*36*). Alternatively, symptomatic 229E infection leads to an increase in nasal mucosal temperature (*35*), which may affect the viscosity of nasal secretions and thereby the production of respiratory droplets.

One limitation of our study is the lack of comorbidities and severe disease manifestations in the rhesus macaque model. However, the age range of the animals used in this study (age range 2-16 years old) is probably a fair reflection of the population currently exhibiting the highest prevalence of SARS-CoV-2 infections, which appears to be younger than early during the COVID-19 pandemic (*37*). Despite the lower pathogenicity of B.1.351 overall, severe disease is still expected to occur in individuals with comorbidities. More targeted epidemiological studies are needed to clarify the impact of variants of concern on the disease burden in various subpopulations with and without comorbidities, while excluding the effect of confounding factors such as the overall number of cases and changes in human behavior.

The B.1.351 lineage has been associated with reduced neutralization by convalescent sera of humans previously infected with SARS-CoV-2 and in people vaccinated against SARS-CoV-2 (*38*). In addition, reduced vaccine efficacy against mild-to-moderate COVID-19 was observed with the B.1.351 variant for vaccines tested in multi-center clinical trials in South Africa and Qatar (*39, 40*). Rechallenge studies in hamsters suggest that prior exposure with SARS-CoV-2 protects against disease but not re-infection of the upper respiratory tract (*41*). These studies, together with our side-by-side comparison of three SARS-CoV-2 variants in the most relevant animal model for preclinical development and comparative pathogenicity, suggest that ongoing circulation under diverse evolutionary pressures favors transmissibility and immune evasion rather than an increase in intrinsic pathogenicity.

## Acknowledgements

We thank Tina Thomas, Rebecca Rosenke and Dan Long for assistance with histology; RMVB animal care staff for taking care of the animals, Jon Schulz, Elaine Haddock, Stacey Rickleffs, Kent Barbian, Kathy Cordova, Marissa Woods and Carl Shaia for technical assistance; Anita Mora for assistance with figures, Mukul Ranjan and Kimberly Stemple for assistance in acquiring the VOC isolates, and Sujatha Rashid at BEI Resources for advice on culturing conditions for the VOC isolates. We thank Cara Bushmaker (Marcus Daily Memorial Hospital) for providing the clinical specimen resulting in the D614G isolate used here, and Andrew Pekosz at Johns Hopkins University Bloomberg School of Public Health for the B.1.351 isolate. SARS-CoV-2 isolate hCoV-19/England/204820464/20200, NR-54000, contributed by Bassam Hallis, was obtained through BEI Resources, NIAID, NIH.

## Funding

This study was supported by the Intramural Research Program of NIAID, NIH.

## Author contributions

Conceptualization, V.J.M. and E.d.W.; investigation, V.J.M, M.F., M.S., B.N.W., F.F., L.P.-P., B.B., M.G.H., D.R.A., A.O., P.W.H., B.J.S., J.L., S.L.A., C.M., N.v.D, G.S. and E.d.W.; writing-original draft, V.J.M. and E.d.W.; writing, review and editing, all authors.

### Competing interests

The authors have no conflicts of interest to declare.

## Supplemental materials

**Table S1.**
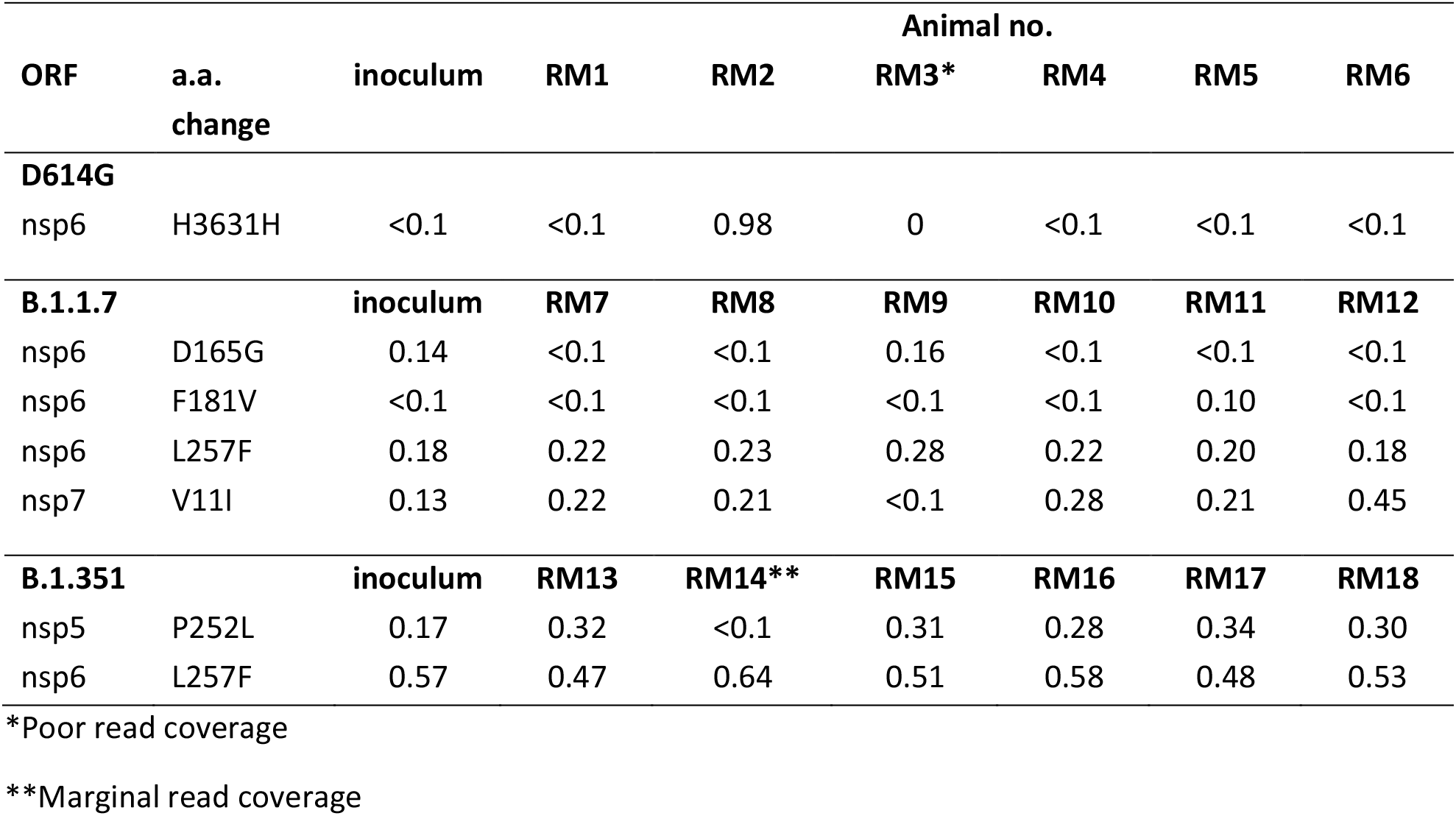
Genome changes observed in virus inoculum and 2dpi BAL samples as compared to reference sequence. Changes are indicated in allelic fraction calculated after filtering as indicated in Methods.

**Figure S1.**
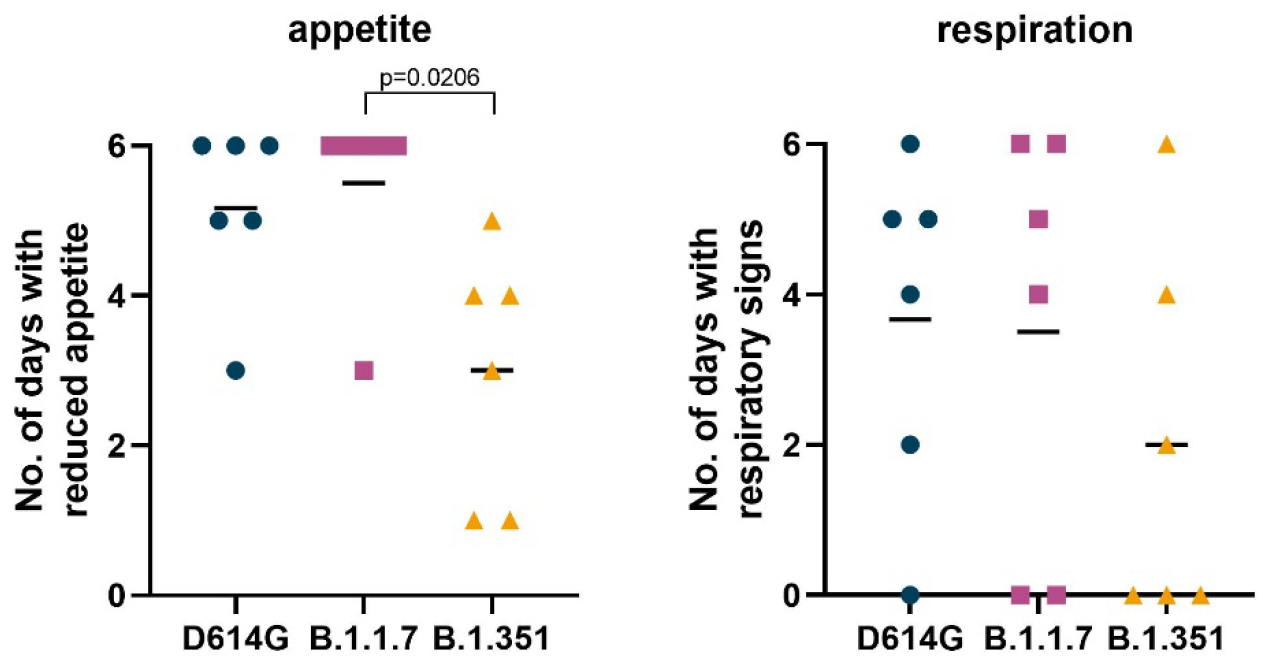
Clinical disease observed in rhesus macaques inoculated with D614G, B.1.1.7 and B.1.351. Three groups of six adult rhesus macaques were inoculated with SARS-CoV-2 variants D614G, B.1.1.7 or B.1.351. After inoculation, animals were observed for disease signs and scored according to a pre-established clinical scoring sheet. The number of days an animal showed reduced appetite (left panel) or changes in respiration pattern (right panel) are indicated. Statistical analysis was performed using a Kruskal-Wallis test with Dunn’s multiple comparisons; p-values <0.05 are indicated.

**Figure S2.**
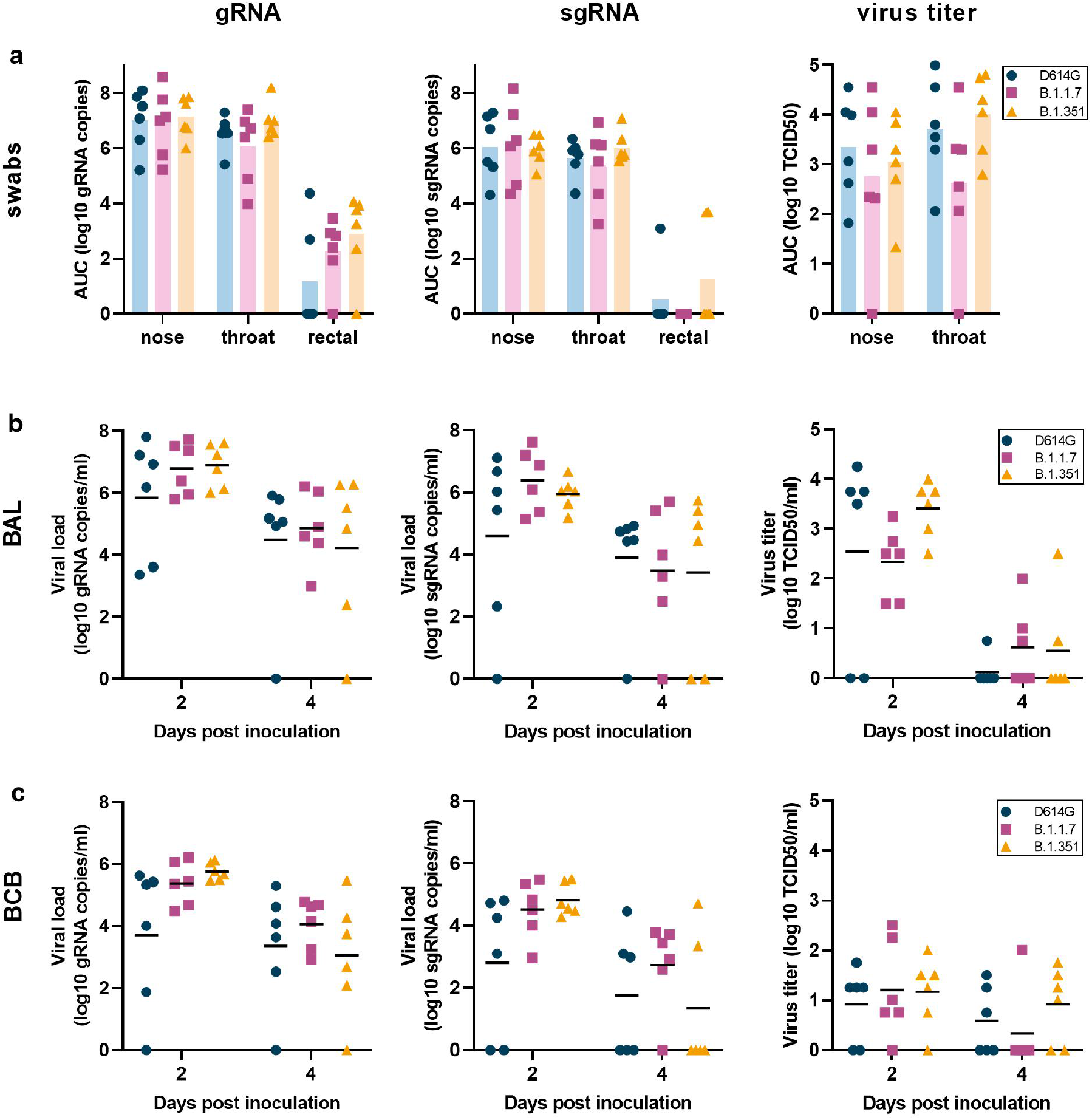
SARS-CoV-2 viral loads and virus titers in swabs, bronchoalveolar lavages and bronchial cytology brushes. Three groups of six adult rhesus macaques were inoculated with SARS-CoV-2 variants D614G, B.1.1.7 or B.1.351. After inoculation, clinical exams were performed on 2, 4, and 6 dpi during which nose, throat, and rectal swabs were collected. qRT-PCR was performed to detect genomic (left column) and subgenomic RNA (middle column), and virus titration was performed to detect levels of infectious virus (right column) and the Area Under the Curve (AUC) was calculated as an indication of the total amount of virus shed in these samples. Swabs (a), bronchoalveolar lavages (BAL) (b) and bronchial cytology brush (BCB) samples (c) were collected on 2 and 4 dpi and analyzed for the presence of gRNA, sgRNA and infectious virus. Lines indicate the mean. Bars (a) and lines (b, c) indicate the mean. Statistical analysis was performed using a Kruskal-Wallis test with Dunn’s multiple comparisons tests (a) or a 2-way ANOVA with Tukey’s multiple comparisons test (b, c); no p-values <0.05 were found.

**Figure S3.**
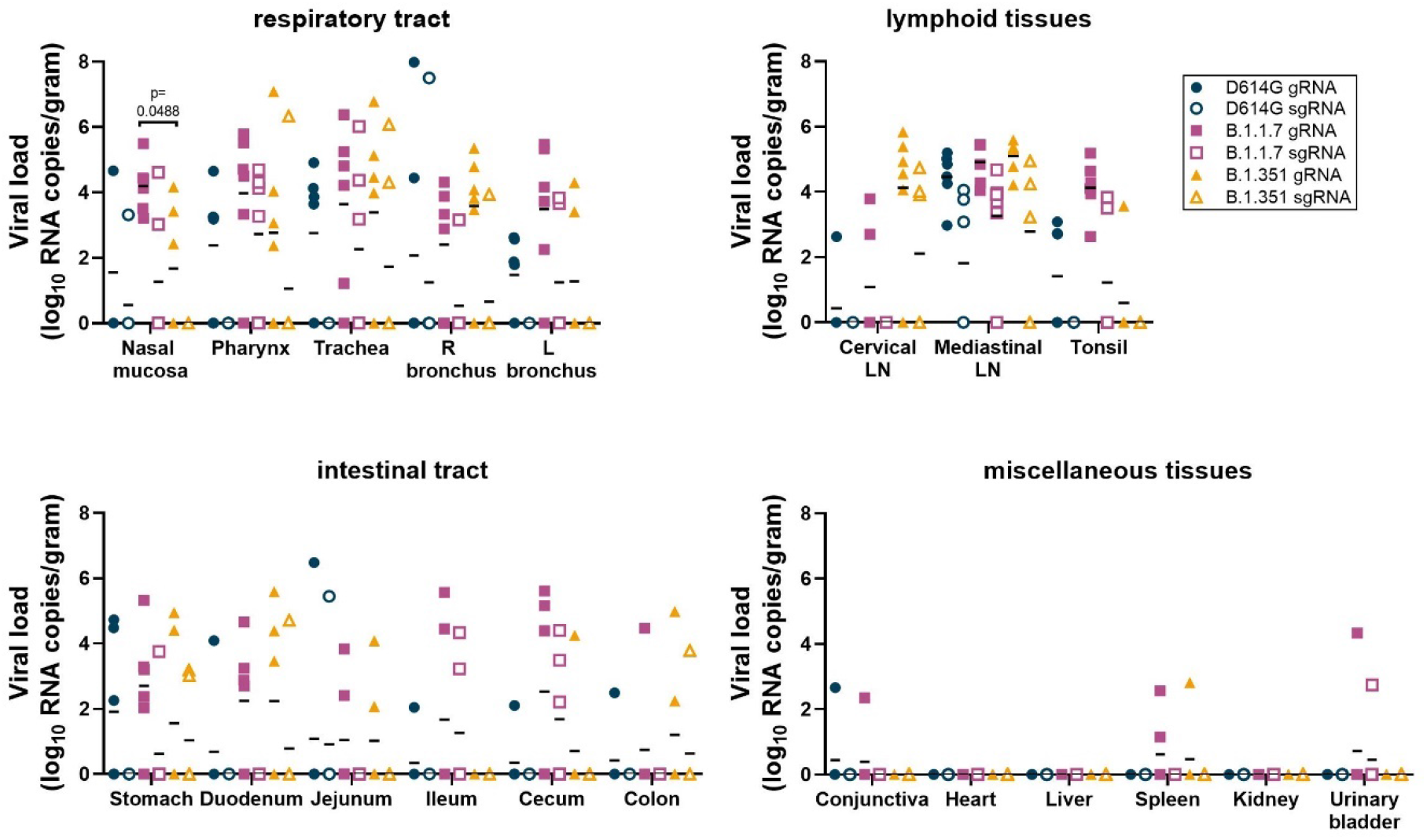
Viral loads in tissues collected from D614G, B.1.1.7 or B.1.351-inoculated rhesus macaques on 6 dpi. Three groups of six adult rhesus macaques were inoculated with SARS-CoV-2 variants D614G, B.1.1.7 or B.1.351. On 6 dpi, all animals were euthanized and necropsies were performed. Samples were collected from many different organs and analyzed for the presence of gRNA (closed symbols) and sgRNA (open symbols). Lines indicate the mean. Statistical analysis was performed using a 2-way ANOVA with Tukey’s multiple comparisons test; p-values <0.05 are indicated.

**Figure S4.**
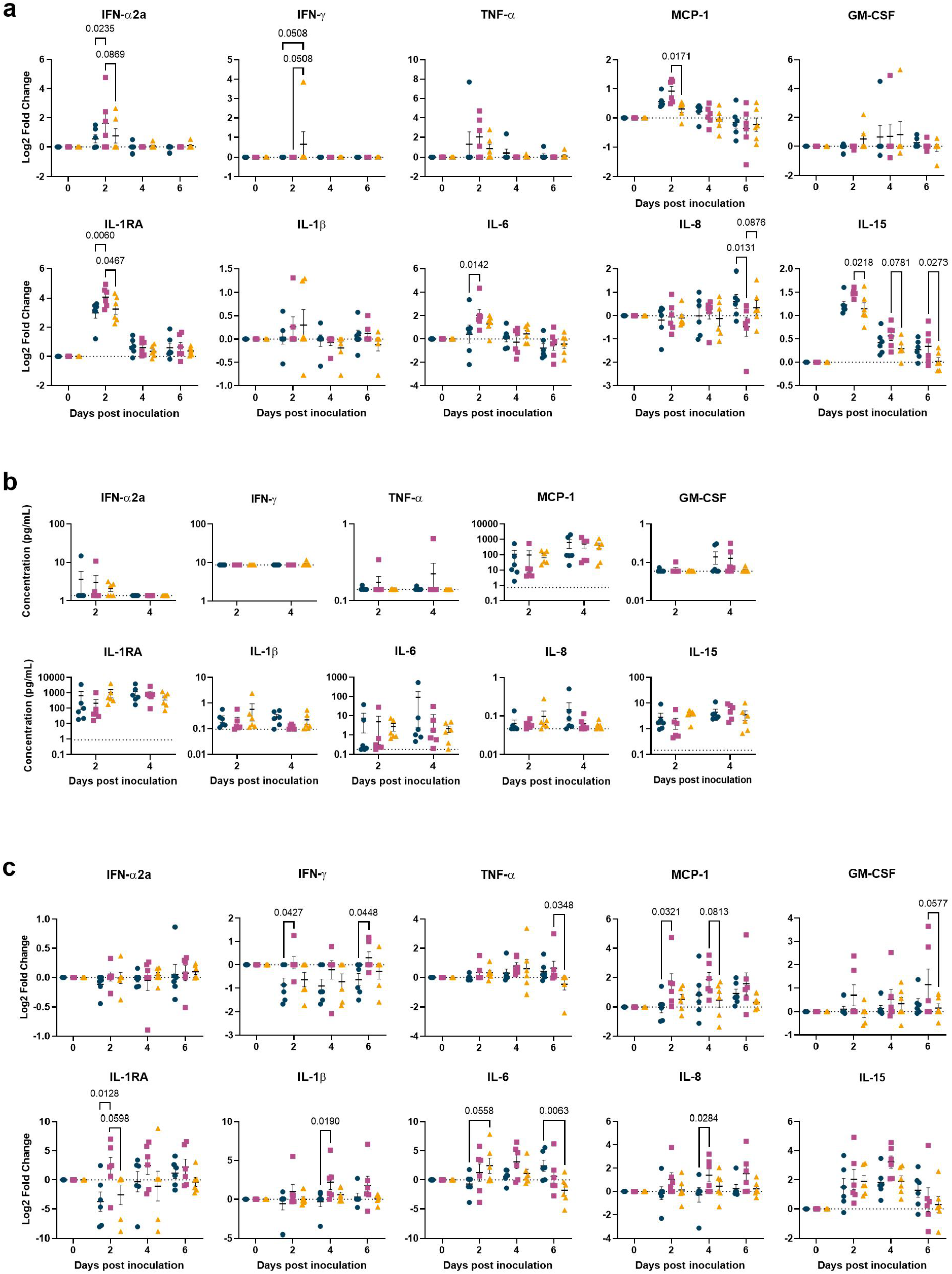
Cytokine and chemokine responses in rhesus macaques inoculated with a D614G, B.1.1.7 or B.1.351 isolate of SARS-CoV-2. The concentration of 10 different cytokines and chemokines were measured in serum (a), BAL (b), and nasal samples (c) at different timepoints before and after inoculation. Fold 2 log changes were calculated for samples where baseline (0 dpi) values were available (a, c); in the absence of a baseline sample concentrations are plotted in pg/mL (b). Blue circles: D614G; pink squares: B.1.1.7; yellow triangles: B.1.351. Dotted lines indicate no change from baseline (a, c) or the maximum lower limit of detection calculated across plates (b). Statistical analysis was performed using a 2-way ANOVA with Tukey’s multiple comparisons test; p-values <0.05 are indicated.

## Notes

### Competing Interest Statement

The authors have declared no competing interest.

